# N-myristoyltransferase: Tracing Steps Backwards to Find a Way Forward

**DOI:** 10.1101/2020.10.23.352898

**Authors:** Dean Reddick, Daniel I Udenwobele, David Datzkiw, Revanti Mukherjee, Shailly Varma Shrivastav, Sara Good, Anuraag Shrivastav

**Affiliations:** Department of Biology, University of Winnipeg, Winnipeg, Manitoba, Canada; Oncodrex Pharmaceuticals, Winnipeg, Manitoba, Canada; Research Institute of Oncology and Hematology, CancerCare Manitoba, Winnipeg, Manitoba, Canada

## Abstract

N-myristoylation refers to the attachment of a 14-carbon fatty acid onto the N-terminal glycine residue of a target protein. The myristoylation reaction, catalyzed by N-myristoyltrasnferase (NMT), is essential for regulating cellular activities such as signal transduction, proliferation, migration, differentiation, and transformation. Although a considerable amount of research is performed on the overexpression of NMT in pathogenic conditions, a fundamental knowledge gap exists on the evolution of NMT and the functional impact of myristoylation for normal cellular development and functions. We performed evolutionary analyses of the NMT gene and found that most non-vertebrates harbor a single nmt gene and all vertebrates examined harbor two genes; nmt1 and nmt2. For the first time, we report that teleosts harbor two copies of nmt1, named nmt1a and nmt1b. We traced the evolutionary history of the chromosomal fragments hosting NMT1 and NMT2 in humans and found that NMT1 and NMT2 trace back to a single vertebrate ancestral chromosome. We also report the presence of putative nuclear localization sequence (NLS) and amino acid residues flanking NLS. The presence of phosphorylatable amino acid residues flanking the NLS suggests that nuclear localization of NMT is regulated by phosphorylation. The nuclear localization of NMT suggest its potential role in gene transcription.

## Introduction

N-myristoyltransferase (NMT) catalyzes myristoylation reactions. Myristoylation is described as the covalent attachment of a 14-carbon saturated fatty acid onto an N-terminal glycine or lysine residue [1] of a nascent polypeptide. The reaction follows the removal of an N-terminal methionine residue by methionine aminopeptidase [2]. NMT has been found to occur in a wide variety of organisms, including mammals [3], plants [4], nematodes [5], fungi [6], and protozoans [7]. In general, myristoylation of a protein will aid in subcellular membrane localization by increasing the protein’s hydrophobicity. This allows the newly myristoylated protein to insert itself securely into the interior, hydrophobic region of a membrane, including the plasma membrane and subcellular membranes such as the endoplasmic reticulum and the nuclear membrane [8]. Thus, myristoylation is a crucial process for some proteins enabling them to localize to their intended destination and function. Myristoylated proteins are found in diverse physiological activities such as cell signalling, signal transduction, cellular differentiation, and cellular transformation [9–11]. Myristoylation has also been shown to stabilize proteins and facilitate protein-protein interactions [12].

In mammals and higher vertebrates, N-Myristoyltransferase exists as two distinct but structurally similar enzymes, NMT1 and NMT2. Both forms of the enzyme catalyze the irreversible attachment of a myristic acid onto a target protein. Two genes, Nmt1 and Nmt2, encode the NMT1 and NMT2 proteins, respectively. NMT1 and NMT2 share approximately 77% amino-acid homology [13], with a highly conserved C-terminal catalytic region and the N-terminal region with substantial sequence variation [14]. Although the two enzymes share similar substrates, biochemical and kinetic studies indicate that they are not functionally redundant. For instance, previously, Nmt1 -/- (homozygous knockout) mice died between embryo development days 3.5 and 7.5, suggesting that Nmt1 was essential for early embryo development. Northern blots revealed lower levels of Nmt2 expression in early development than at later time points, a potential explanation for the demise of Nmt1−/− embryos. In Nmt1−/− embryonic stem (ES) cells, Nmt2 mRNA was present, but the total NMT activity levels were reduced by ~95%, suggesting that Nmt2 contributes little to the total enzymatic activity in these early embryonic cells [15]. We also demonstrated that NMT1 is essential for the proper development of myeloid lineage by knockout experiments wherein NMT2 could not rescue the aberrant developmental process. These reports confirmed that NMT1 and NMT2 are not redundant in function, although they both catalyze the myristoylation reaction. Another study also demonstrated differences between the two enzymes (ducker ref). The knockdown of Nmt1 inhibited cell replication associated with the loss of activation of c-Src and its effector protein - focal adhesion kinase (FAK), while knockdown of Nmt2 gene did not.

Further, the depletion of either NMT1 or NMT2 induced apoptosis, but NMT2 depletion showed a 2.5-fold greater effect than NMT1. Additionally, knockdown of NMT2 expression resulted in the induction of apoptosis by the BCL family of proteins, whereas knockdown of NMT1 did not. Finally, intramural injection of Nmt-specific siRNA for both Nmt1 and Nmt2 together reduced tumor growth, while injection of Nmt2-specific siRNA alone failed to reduce tumor growth [16].

In humans, both NMT forms are expressed in most of the body’s tissues, and their dysregulation is implicated in the development and progression of several diseases [17, 18]. For instance, NMT activity and expression has previously been reported to be upregulated in adenocarcinomas, hormone-receptor-positive breast cancers, oral squamous cell carcinomas, brain tumors, and gallbladder carcinomas [19, 20]. Clegg et al. reported that the cell proliferation rates were strongly correlated with NMT expression in mammary epithelial cells and described the redistribution of NMT from the nuclear membrane to the cytosol in cancer cells [21]. Recent studies have shown that in-vitro inhibition of NMT slows the progression of viral diseases such as the common cold [22] and HIV [23].

The involvement of NMT in various metabolic diseases such as cancer and diabetes, involvement in infectious diseases warrants more in-depth insight into the evolution of NMT gene(s) to understand and predict functional implication of NMT in pathogenic conditions. Therefore, we set out to perform detailed phylogenetic and evolutionary analyses employing bioinformatics tools. We report a highly conserved nuclear localization signal (NLS) present within the primary structure of NMT that may be responsible for the occasional localization of NMT in the nuclei of cells.

## Methods

### NMT Multiple Sequence Alignment and Phylogenetic Analysis

Sequence collection: NMT protein sequences were collected from all branches of the eukaryotic tree, including unicellular eukaryotes from the stramenopiles (or heterokonts, (e.g., Phaeodactylum tricornutum and Phytophthora infestans), Alveolata (Plasmodium falciparum -Apicomplexa, Tetrahymena thermophila – Ciliates), and Rhizaria (e.g., Bigelowiella natans) – also known as the SAR supergroup, representing a diverse group of distantly related protozoans. Additionally, we included sequences of the Amoebozoa (e.g., Dictyostelium purpureum, Acytostelium subglobosum), Excavates (e.g., Leishmania major – Euglenoza), Algae (e.g., Crypophytes - Guillardia theta, e.g., Chlorophyta - Chlamydomonas reinhardtii) and its’ sister group Land Plants (including representatives of liverworts, bryophytes, and flowering plants). Finally, we sampled more extensively from the Opisthokonta supergroup including Fungi (e.g. Schizosaccharomyces pombe, Saccharomyces cerevisiae), Apusomonads (e.g. Thecamonas trahens), choanoflagellates (Monosiga brevicollis) as well as animals including Poriphera (Sponges, e.g. Amphimedon queenslandica), diverse protostomes (including nematodes, annelids, amd molluscs) and most broadly from the deuterostomes - including basal taxa Strongylocentrotus purpuratus (echinoderm) and Branchiostoma floridae (cephalochordates), and then within chordates – sea lamprey (Agnatha), elephant shark (Chondrichythes), spotted gar (a pre-3R fish) and representatives of both teleosts and tetrapods (see SuppFile_AccessionNumbers.xls for a full list of species and accession numbers). Sequences were collected from Ensembl genomes (using the Ensembl metazoa, animals, plants, and fungi) using the Ensembl gene trees to identify nmt sequences from diverse taxa and NCBI using a blastp for the following taxa; birds, amphibia, teleosts, echinoderms, hemichordates, and cephalochordates.

Phylogenetic reconstruction: Sequences with less than 200 amino acids or greater than 500 amino acids were not retained for further analyses. Two alignments were generated, one for the pan-eukaryotic analyses and one for deuterostome-specific analyses. The N-terminus of NMT is highly variable in length and sequence, which is problematic for global multiple sequence alignments (MSA), especially at the pan-eukaryotic level. Thus, alignments were performed using the e-INS algorithm and MAFFT-DASH to guide the alignment with structural information from homologous proteins [24]. Aligned sequences were trimmed to remove regions for which amino acids were present in only a single taxon. The best model of amino acid substitution was estimated by maximum likelihood (ML) and implemented to reconstruct the relationship among sequences using ML with 500 bootstraps replicates in the program MegaX [25]. The phylogenetic tree and alignment were plotted using the R library ggtree [26].

Origin of vertebrate nmt1 and nmt2. We followed the approach of Yegorov and Good [27] and Sacerdote et al [28] to trace the evolutionary history of nmt1 and nmt2 in vertebrates. Briefly, this involves tracing the genomic fragments housing nmt1 and nmt2 genes across vertebrate taxa to the ancestral fragments on which they were housed in the pre 1R and pre 3R vertebrate genomes to examine whether a) nmt1 and nmt2 in humans were derived from a single gene in the pre 1R genome and b) teleost specific duplicates nmt1a and nmt1b emanate from a single gene in the pre 3R fish genome.

### Nuclear Localization Signal (NLS) Identification

NLS Mapper (http://nls-mapper.iab.keio.ac.jp/cgi-bin/NLS_Mapper_form.cgi) and NLStradmus software (http://www.moseslab.csb.utoronto.ca/NLStradamus/) were used to analyze the putative NLS’s in the primary structures of human NMT1 (hNMT1) and hNMT2. NucPred software (https://nucpred.bioinfo.se/nucpred/) was used to predict whether hNMT could reside in the nucleus or not.The NLStradums software includes three models that can be used to predict an NLS in the queried protein. A 2-state static Hidden Markov Model (HMM), a 2-state dynamic HMM, and a 4-state static HMM. The 2-state models are designed to predict classical monopartite NLS’s while the 4-state is designed to predict bipartite NLS’s [29]. For this study, FASTA formatted hNMT primary structure sequences were uploaded into the NLStradmus software, and the 2-state static model was selected for NLS prediction. The software also includes a posterior cut-off threshold score. The higher the threshold used, the better the predictive power of the software. The maximum score which can be selected is 1.0. The NLStradmus software recommends using a cut-off threshold value of no lower than 0.5 (31). A threshold of 0.7 and 0.8 was used to predict the NLS of NMT1 and NMT2, respectively.

NLS mapper accurately predicts NLSs specific to the importin αβ pathway by calculating NLS activity levels using four NLS profiles; class 1/2, class 3, class 4, and bipartite NLSs. These profiles were generated by an extensive amino acid replacement analysis for each NLS class in budding yeast. NLS mapper extracts putative NLS sequences with a score equal to or more than a user-selected cut-off score. Briefly, an NLS with a score of 8 - 10 indicates exclusive nuclear localization. A score of 7 or 8 indicates partially nuclear localization, 3 - 5 indicates both the nuclear and cytoplasmic localization whereas, a score of 1 or 2 is for cytoplasmic localization [30]. The user must also select the region of the uploaded sequence to be analyzed. Specifically, the analysis can be restricted to the first 60 amino acids of the protein (N-terminus) or expanded to include the entire primary sequence. For this study, the primary structures of hNMT1 and hNMT2 were uploaded into the NLS mapper software in FASTA format. A cut-off score of 4 and 6 were selected for NMT1 and NMT2, respectively. The analysis was unrestricted to include the entire primary structure of NMT1 but was restricted to include only the first 60 amino acids of NMT2.

NucPred software analyzes patterns in eukaryotic protein sequences and predicts if a protein spends at least some time in the nucleus or no time at all. The program applies genetic programming (GP), a machine-learning technique that automatically develops computer programs in an artificial evolutionary process. The evolved predictors incorporate multiple regular expressions that are matched against the input (amino acid) sequence. For single sequences, NucPred calculates a per-residue scoring. Each amino acid location is colored according to its influence on a ‘nuclear’ classification. The closer the color of a sub-sequence lies at the red end of the color spectrum, the more positive is the effect. The program also provides a NucPred score, reflecting the probability of the uploaded protein residing in the nucleus. The higher the score, the more likely the protein will occupy the nucleus. The maximum score attainable is 10 [31]. For this study, FASTA formatted hNMT1 and hNMT2 primary sequences were submitted in the NucPred software.

## Results

### NMT Multiple Sequence Alignment Analysis

Pan-eukaryotic nmt phylogeny. Reconstruction of the phylogenetic relationship among eukaryotic taxa indicates that all branches of the eukaryotic tree harbor nmt (Figure 1), and the phylogenetic placement of most taxa recapitulated their taxonomic position (please see Supplementary file for the tree with species names). Only a single nmt sequence was found in all lineages except deuterostomes (SuppFile_ACCESSION NUMBERS) for most taxa. The N-terminus of the gene is highly variable among taxa, and for species for which the sequence was based on ab-initio prediction without expression data, it may be missing from the alignment (SuppFileAccessionNumbers). Nevertheless, one salient feature of the N-terminal end is that animals and some fungi harbour a putatitve nuclear localization sequence (NLS), which is not present in any of the sequences of unicellular eukaryotes, nor in floweing plants (see SuppFile_Eukaryoticalignment.fas). For fungi, which are in the supergroup Opisthokonta with animalia, some taxa harbor an NLS motif similar to that in animals (e.g., Neurospora crassa sequence is “KKKNKKKSKK”), while other fungi did not possess this motif but have other motifs – such as “NNKNTKNSQQ” in Schizosaccaharomyces_pombe. On the other hand, all animals had an NLS motif situated in a homologous region of the mature peptide (Figure 1).

**Figure 1.**
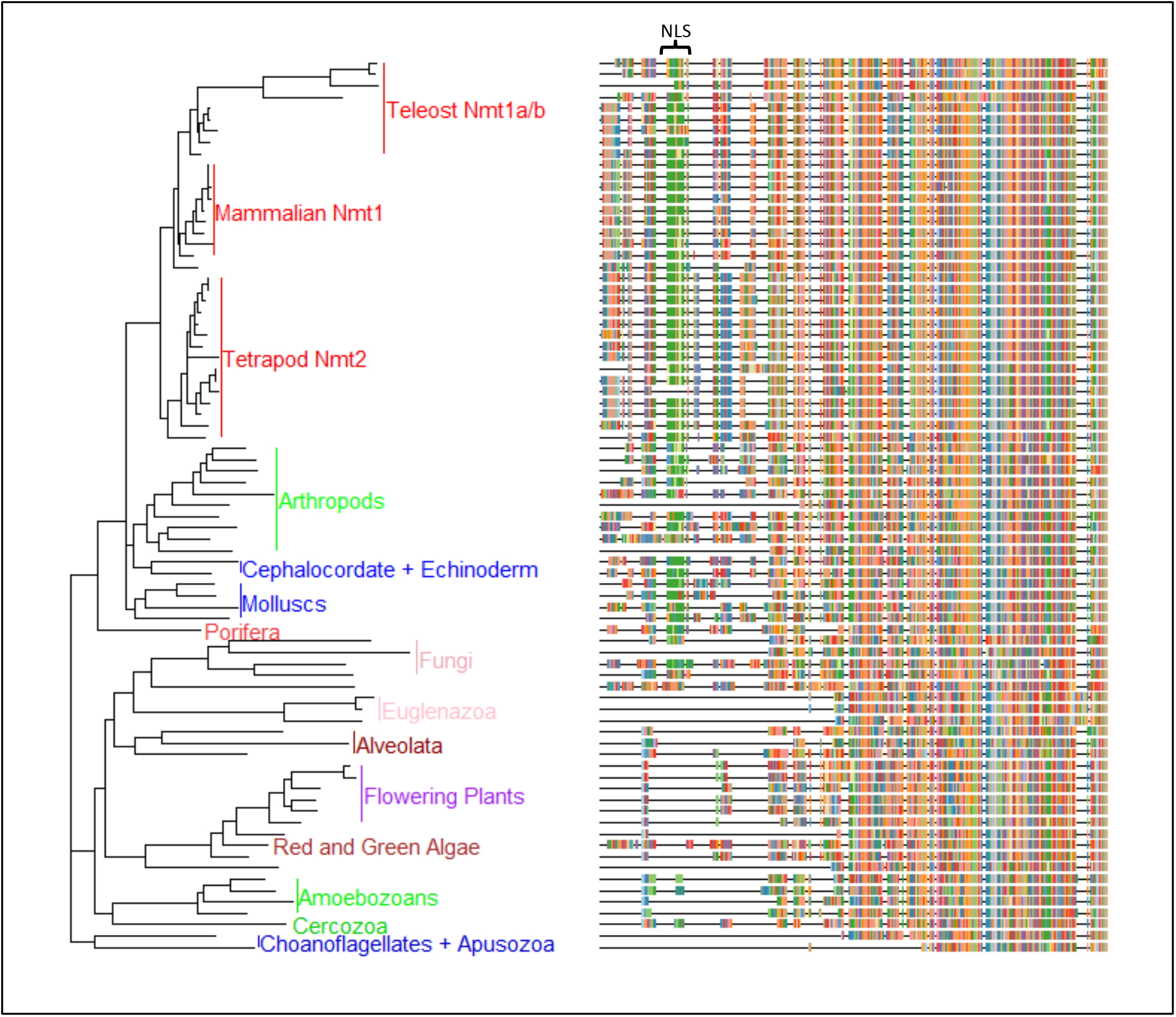
Evolutionary history of NMT sequences across eukaryotes as inferred by Maximum Likelihood method and Le_Gascuel_2008 model of protein evolution (44). The tree with the highest log likelihood (−31889.84) is shown. Initial tree(s) for the heuristic search were obtained automatically by applying Neighbor-Join and BioNJ algorithms to a matrix of pairwise distances estimated using a JTT model, and then selecting the topology with superior log likelihood value. A discrete Gamma distribution was used to model evolutionary rate differences among sites (6 categories (+G, parameter = 0.7512)). The tree is drawn to scale, with branch lengths measured in the corrected number of substitutions per site. This analysis involved 79 taxa with a total of 595 amino acid positions in the final dataset. Evolutionary analyses were conducted in MEGA X (28).

### Phylogenetic relationship of NMT1 and NMT2 in vertebrates

While most non-vertebrates harbor a single nmt, all vertebrates examined harbor two genes; nmt1 and nmt2, and teleosts additionally harbor two copies of nmt1, named nmt1a and nmt1b. By tracing the evolutionary history of the chromosomal fragments hosting NMT1 and NMT2 in humans (CHr17, 45.051Mb, and Chr10, 15.102 Mb respectively), we identified that NMT1 and NMT2 trace to a single vertebrate ancestral chromosome, VAC “E” based on Nakatani et al., or to chromosome 1 of the pre-1R vertebrate genome as reconstructed by Sacerdote et al., [28]. This indicates that NMT1 and NMT2 are derived from a single ancestral vertebrate gene and are “ohnologues.” However, most teleosts harbor three genes: two copies of NMT1 (NMT1a and NMT1b) and a single copy of NMT2. The NMT1a/NMT1b paralogues also appear to have duplicated during the fish-specific WGD event. For example, NMT1a and NMT1b are on sections of chromosomes 8 and 19 in medaka that arose from a single ancestral pre-3R Osteichythan chromosome i (see 29 for methodological details), that was duplicated in the neopterygii ~ 330mya. These results are supported by inferences at genomicus, a synteny database (www.genomicus.biologie.ens.fr): choosing NMT1 in human as the focal species and chordates as the ancestor, NMT2 originates in euteleostomi post-2R genome and nmt1a and nmt1b originate in the neopterygii whole genome duplication event. This indicates that a single nmt gene in pre-vertebrates, duplicated to give rise to nmt1 and nmt2, early in vertebrate evolution, and that teleost nmt1a/nmt1b are 3R paralogs. The teleost gene nmt1a gene shares high amino acid similarity to mammalian nmt1, but the teleost Nmt1b gene is divergent, and groups basal to both genes in the vertebrate phylogeny (Figure 2).

**Figure 2.**
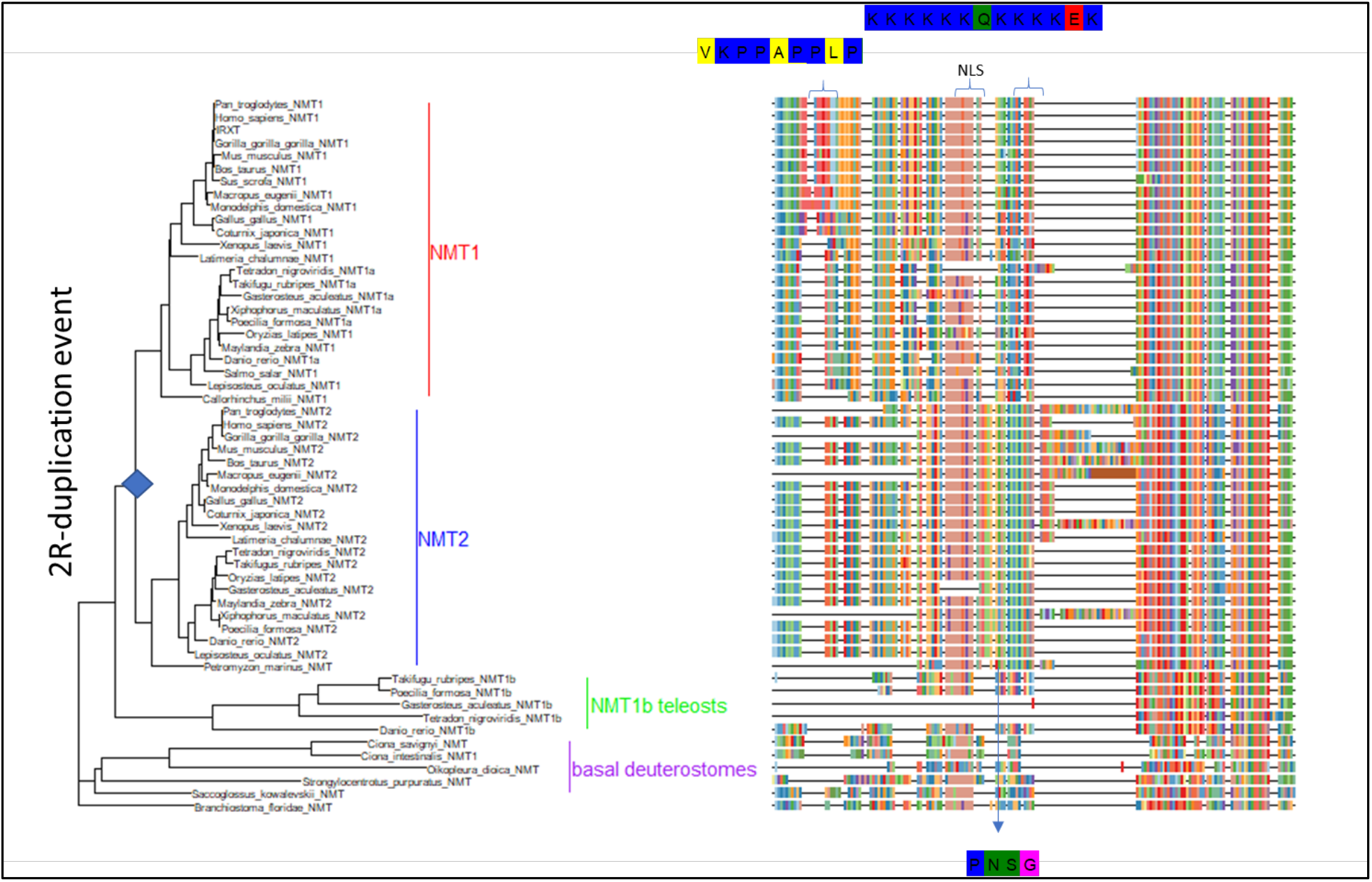
Evolutionary history of deuterostome NMT sequences inferred using the Maximum Likelihood method and Jones et al., model of sequence evolution (45), with 500 bootstraps. The tree with the highest log likelihood (−14416.10) is shown. Initial tree(s) for the heuristic search were obtained by applying Neighbor-Join and BioNJ algorithms to a matrix of pairwise distances estimated using a JTT model, and then selecting the topology with superior log likelihood value. A discrete Gamma distribution was used to model evolutionary rate differences among sites (6 categories (+G, parameter = 0.6435)). The tree is drawn to scale, with branch lengths measured in the number of substitutions per site, given a total of 595 amino acids. The evolutionary analyses were conducted in MEGA X (45); the tree with bootstrap values is shown in SuppFile_VertTreeBoot.png. On the right, the first 150 amino acids of the alignment for each of the taxa in the tree is given, the full alignment is shown in SuppFile_VertAlignment.fas.

### Nuclear Localization Signal (NLS)

Two types of NLS have are present in eukaryotic proteins. The classical and non-classical NLS. The classical version possesses several basic amino acids such as lysine, glutamine and arginine residues [32]. Classical NLS’s can be further subdivided into a monopartite and bipartite type. Monopartite NLS’s possess several basic amino acids in an unbroken sequence. Bipartite NLS’s also contain basic amino acids, but they typically exist as two short clusters separated by a spacer sequence [33]. For most species included in the alignment, the sequence contains several basic amino acids, including nine lysine, one glutamine and one arginine residues. Such an NLS would allow NMT to interact with importin-α and be transported through a nuclear membrane pore into the nucleus through the classical nuclear localization pathway [34].

A closer inspection of the alignment affirms that the NLS of NMT is flanked by several highly conserved serine, threonine and tyrosine residues. For this study, residues between positions 30 - 90 were considered to be flanking the NLS. For NMT1, these include S31, S40, Y41, S47, T52, S69, T71, S73, and S83. For NMT2, they include T32, S38, T60, S61, S64, S66, S68, and S74. It is plausible that any or all these highly conserved residues serve as phosphorylation sites on the NMT enzymes regulating the tertiary arrangement of the NLS and thus regulating the subcellular localization of NMT [35].

### NLS in NMT1 and NMT2

NLS mapper and NLStradmus software were used to investigate the highly conserved poly lysine sequence of NMT1 and NMT2. The NLStradumus software predicted the poly lysine sequence of hNMT1 and hNMT2 as promising candidates for monopartite NLS’s at a cut-off value of 0.7 for NMT1 and a cut-off value of 0.8 for NMT2 (Figure 3). The predicted NLS of hNMT1 is located at position 55 to 67 and 46 to 58 for hNMT2. This correlates with the positions of the polylysine sequence observed in the multiple sequence alignment. The NLS mapper predicted a bipartite NLS for NMT1, which included the same monopartite poly lysine sequence predicted by NLStradmus, along with an additional sequence from residues 298 to 330. A monopartite NLS was predicted for NMT2 from residue 50-58, which is very similar to the prediction made by NLStradmus for NMT2 (Figure 4).

**Figure 3.**
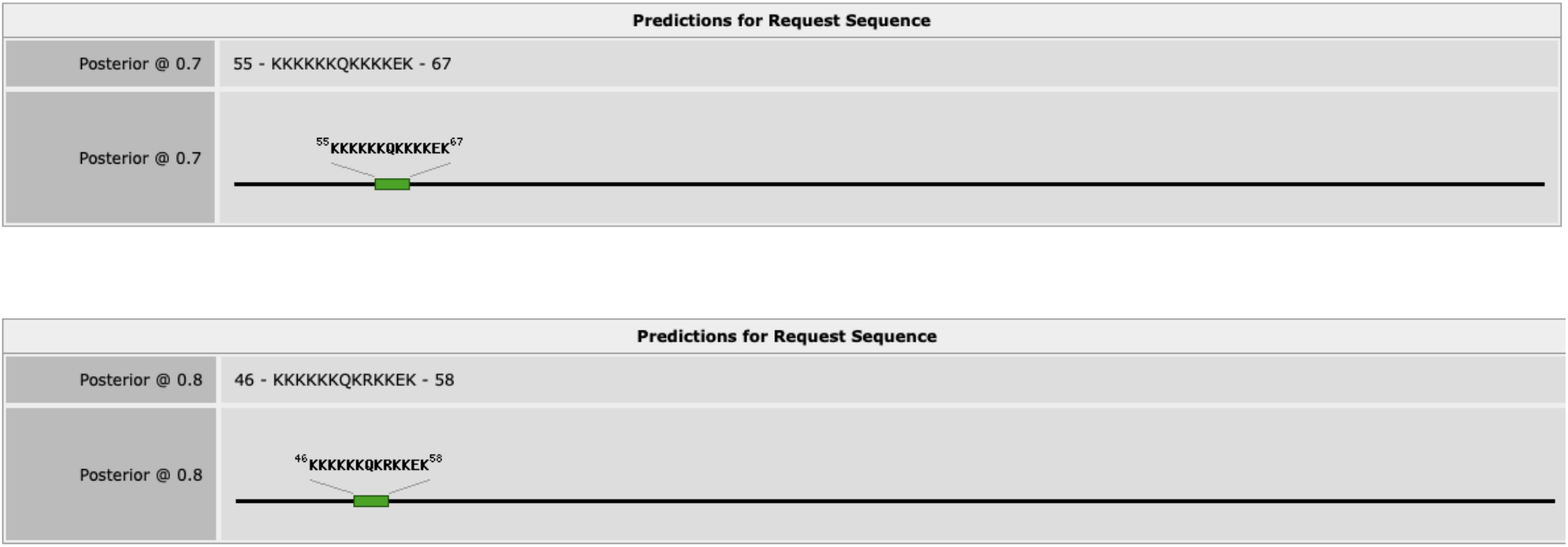
NLStradmus results for the prediction of an NLS in the primary amino acid sequence of hNMT1 (top) and hNMT2 (bottom). The black line in the image represents the entire primary structure of the NMT proteins while the green box represents the relative positioning of the NLS within the primary sequence. A posterior prediction value of 0.7 was selected for NMT1 and 0.8 was selected for NMT2. Based on statistical modeling, NLStradmus recommends staying above a cut-off value of 0.5 for accurate results and higher cut-off value represents a stronger result.

**Figure 4.**
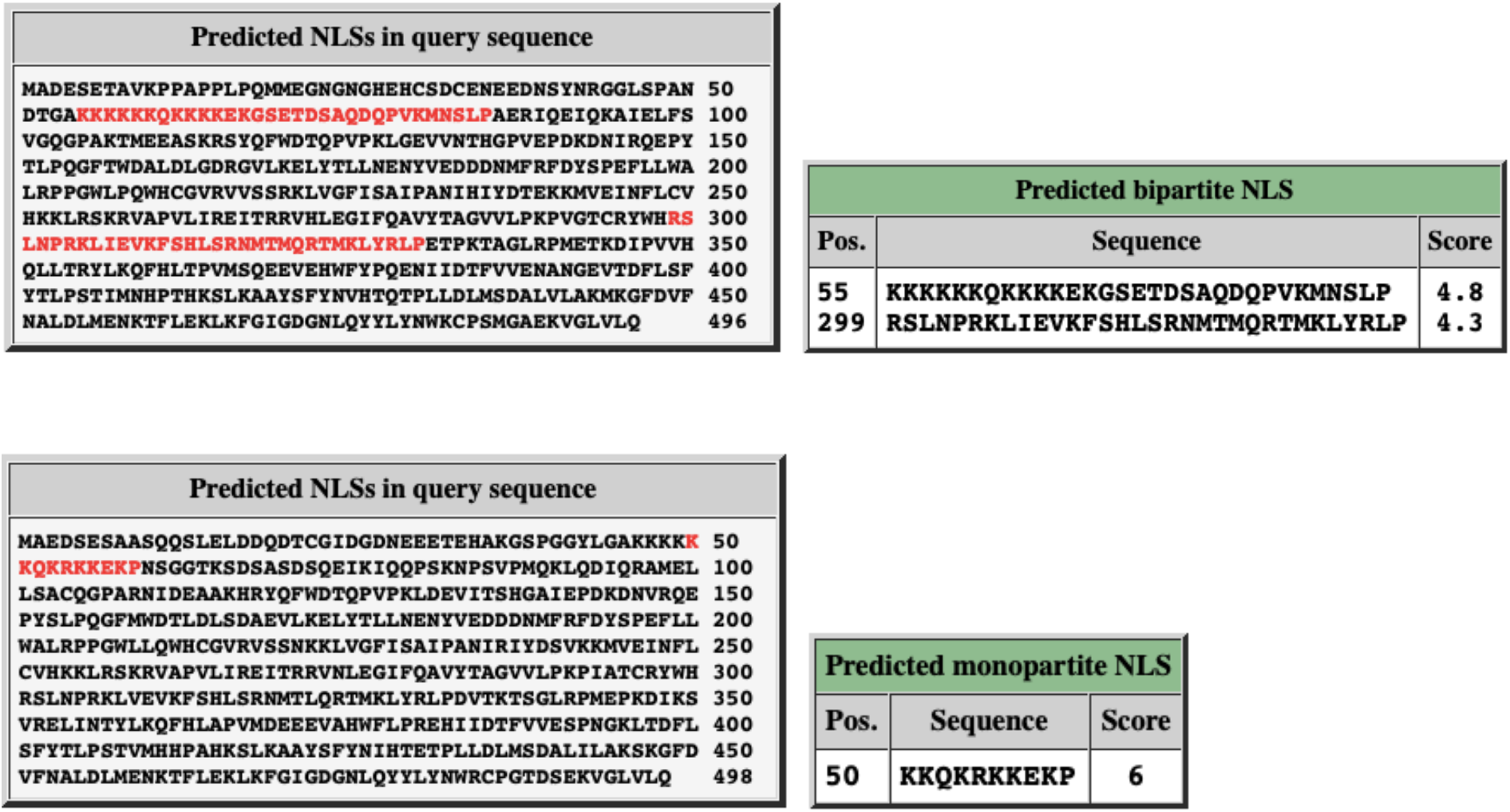
NLS of hNMT1 (top) and hNMT2 (bottom) predicted by NLS Mapper software. A cut-off score of 4.8 was used to predict the poly lysine sequence of NMT1 as an NLS while a cut-off score of 6 was used to predict the poly lysine sequence of NMT2 as an NLS.

Following the putative NLS prediction, it was necessary to determine the probability of NMT proteins residing in eukaryotic cells’ nuclei. To meet this objective NucPred software was employed. For this study, hNMT1 or hNMT2 primary sequences in FASTA format were submitted into the NucPred prediction software. A NucPred score of 0.44 was given for NMT1, and a score of 0.81 was given for NMT2. Partial yellow to light-green coloring of the putative NLS can be seen in the NMT1 sequence, while red, orange, and yellow coloring can be seen in the NLS for NMT2 (Figures 5 and 6).

**Figure 5.**
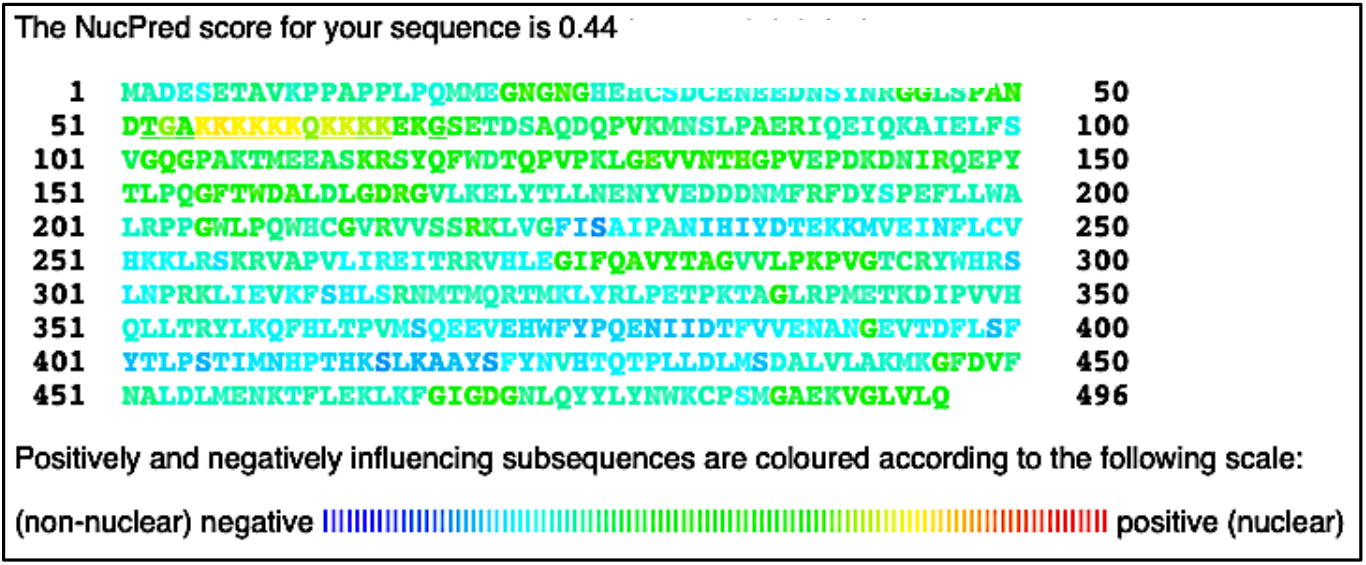
Results of NucPred analysis of hNMT1 primary sequence. Colors toward the red end of the visible spectrum signify amino acids which contribute to nuclear localization of NMT1. Colors toward the blue end signify amino acids that do not contribute to nuclear localization (45). The NMT1 NLS appears yellow and light green in the 51 – 100 residue row.

**Figure 6.**
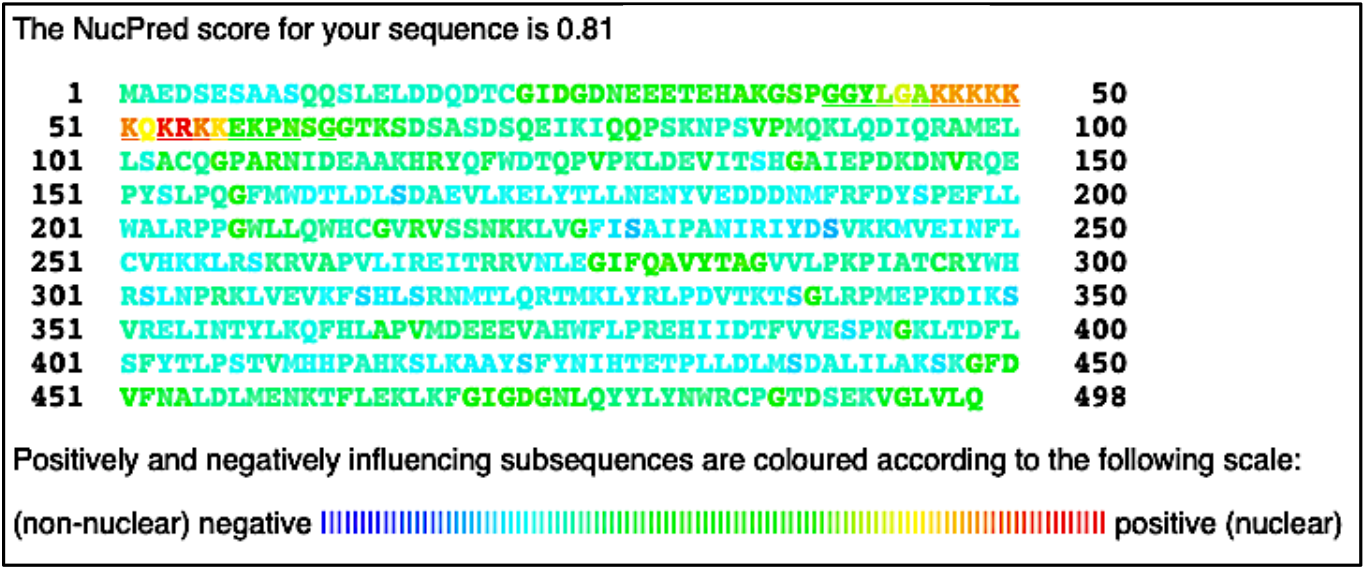
Results of NucPred analysis of hNMT2 primary sequence. Colors toward the red end of the visible spectrum signify amino acids which contribute to nuclear localization of NMT2. Colors toward the blue end signify amino acids that do not contribute to nuclear localization (33). The NMT2 NLS appears red, orange and yellow in the 1 – 50 and 51 – 100 residue rows.

## Discussion

We find that most non-vertebrates harbor a single nmt, while all vertebrates examined have two genes; nmt1 and nmt2, and teleosts additionally harbor two copies of nmt1, named nmt1a and nmt1b. Reconstruction of the phylogenetic relationship among deuterostome NMT sequences indicates that vertebrate NMT2 sequences are monophyletic; the phylogenetic relationship among sequences largely reflects the time of divergence of taxa,, and the single identified lamprey NMT-like sequence is basal to this group (Figure 1). On the other hand, vertebrate NMT1 sequences are more variable in teleosts but are well-conserved in mammals (Figure 1). The highly divergent nature of NMT1b drives this in teleosts: teleost NMT1a clusters as the sister group to mammalian NMT1 sequences, but NMT1b in teleost is divergent and groups basal to both genes in the vertebrate phylogeny (Figure 2). Visualization of the alignment for nmt sequences in vertebrates reveals essential differences between NMT1 and NMT2 in the gene’s N-terminal portion. Notably, mammalian NMT1 harborsharbors an insertion near the N-terminus that is not present in NMT1 in lower vertebrate or fish (Figure 2). In humans, the sequence of this insertion is VKPPAPPLP. Secondly, as observed in the pan-eukaryotic alignment, the NLS for both NMT1 and NMT2 in vertebrates is lysine-rich and highly conserved – most frequently with the motif “KKKKKKQKKKKEK” for NMT1 except in lower vertebrates in which the middle Q is typically P. The NLS sequence motif is very similar, and indeed more conserved for NMT2 than for NMT1, and has the following motif “KKKKKKQKRKKEK.” However, following this motif, NMT2 has an insertion sequence “PN(or S)SG” which is not present for NMT1.

We propose that the NLS of NMTs interact with importin-α shuttling protein to initiate NMT nuclear membrane translocation. Nuclear NMT could interact with and myristoylate neighboring nuclear proteins, including transcription factors, or directly interact with genetic material and play a vitalessential role in gene transcription. The activity, interactions, and subcellular localization patterns of many proteins are regulated by posttranslational modifications such as phosphorylation [36], ubiquitination [37], and acetylation [38]. Of these, phosphorylation is most often utilized by eukaryotic cells, with over 13000 proteins undergoing this process [39]. Furthermore, previous studies have demonstrated that the phosphorylation status of amino acid residues in close proximity to a protein’s NLS can modulate the NLS’s secondary and tertiary structure and regulate the protein’s ability to translocate across the nuclear membrane [40]. we observed that several highly conserved putative phosphorylation sites flank the poly lysine NLS’s of NMT. The phosphorylation status of these sites may contribute to the tertiary structure of NMT and its NLS and regulate the protein’s ability to interact with importin-α shuttling protein and thus play a part in the regulation of NMT subcellular localization.

A nuclear localization signal is a characteristic consensus sequence of amino acids present in the primary structure of a protein, which differentiates the nuclear localizing protein cargo from the remainder of proteins in the cytoplasm. An NLS can be classified into either classical NLS that comprises of basic amino acids, including glutamine, lysine, or arginine. Whereas a non-classical NLS comprises of proline-tyrosine (PY) stretch of residues. Classical NLS can be either monopartite comprising a single stretch of basic amino acids as reported in SV40 large T-antigen [41] or bipartite comprising two basic stretches of amino acids separated by a spacer sequence as observed in Ty1 integrase [42]. The non-classical NLS includes the PY sequence containing motif as reported in the Hrp1 protein found in Saccharomyces cerevisiae [43] and the M9 sequence containing motif as reported in the splicing factor, HnRNP A1. For the first time, we report the presence of classical NLS in the primary structure of NMT, which is putative monopartite. Karyopherin or importins are essential proteins that play a role in the nuclear transport of cargo protein containing the NLS. Protein cargo containing classical NLS can complex with an adaptor protein, importin-α, which in turn can latch onto importin-β that mediates its transport into the nucleus [44]. On the contrary, non-classical NLS can interact directly with importin-β without the involvement of adaptor karyopherin and is transported into the nucleus [45]. The multiple sequence alignment performed affirmed the presence of a highly conserved polylysine sequence in the primary structure of NMT. The sequence alignment displays a polylysine sequence comprising nine lysine, one-glutamine, and one-arginine residues unbroken by any spacer sequence, representing a classical monopartite NLS.

Predictive NLS software also recognized the polylysine sequences of hNMT1 and hNMT2 as monopartite NLS’s. The NLStradmus software predicted the polylysine sequence of NMT1 as an NLS candidate with a confidence score (posterior score) of 0.7 (out of 1.0) and the polylysine sequence of NMT2 with a confidence score of 0.8. NLS Mapper also predicted the polylysine sequences as NLS’s, although the confidence score for NMT1 was lower than expected at 4.8 (out of 10) while the score for NMT2 was better at 6.0. Both NLStradmus and NLS Mapper were able to predict the NLS of NMT2 with higher confidence over NMT1. NucPred software predicted that both hNMT1 and hNMT2, at least partially, does reside within the nucleus. The software gave a confidence score of 0.44 (out of 1.0) for NMT1 and a score of 0.81 for NMT2. Again, predicting that NMT2 is more likely to locate in the nucleus than NMT1. NucPred also color codes the individual amino acids of the queried sequence, including any NLS that is present. Amino acids at the red end of the spectrum are more likely to regulate the protein’s transport into the nucleus. The NLS of NMT1 appears yellow, and light green in the NucPred read out while the NLS of NMT2 appears red and orange. Compared to all surrounding amino acids of the NMT sequences (which all appear dark green and blue), the NLS’s of both NMT1 and NMT2 is likely playing a role in nuclear translocation. However, the NLS of NMT2 appears closer to the red end of the spectrum and likely has a stronger affinity for importin-α. Based on the results obtained from these prediction platforms, it is likely that both hNMT1 and hNMT2, especially NMT2, can be transported inside the nucleus of a cell through the classical importin αβ pathway.

Proteins transported across the nuclear membrane must first be distinguished from the plethora of proteins in the cytoplasm. The NLS of the transported protein must be accessible to importin-α shuttling protein. The tertiary arrangement of the transported protein must expose the NLS not embedded within the protein. The phosphorylation status of residues flanking the NLS is likely to play a role in the tertiary arrangement of the NLS and N-terminal region of NMT. It is plausible that the flanking residues’ phosphorylation status is regulating whether the NLS of NMT is exposed on the surface or embedded within the protein. An exposed NLS would make it possible for Importin to interact with NMT and escort it over the nuclear membrane and into the nucleus. It is crucial to determine which phosphorylated amino acid residues on the NMT protein regulate its nuclear localization.

We found that three different NLS predictive software demonstrated a monopartite NLS in the primary structure of NMT. Monopartite NLS are characteristic of eukaryotic proteins, which are translocated into the nucleus through the importin αβ pathway. Furthermore, predictive phosphorylation databases were used to confirm the presence of phosphorylation sites flanking the NLS of NMT. Such flanking phosphorylation sites have previously been shown to regulate the localization of nuclear eukaryotic proteins. The subcellular localization patterns of proteins can be used to investigate the function of such proteins. NMT in the nucleus may play a crucial role in regulating gene transcription and cellular function.

## Supporting information

NMT1 and NMT2 alignment

Accession numbers

Vertebrate Tree

## Acknowledgement

DR and DD acknowledge CIHR for the Charles Banting Graduate Scholarship. DD and DU acknowledge Queen Elizabeth II Scholarship. RM acknowledges UWSA scholarship and The President’s Award UW. AS acknowledge the funding support from the CancerCare Manitoba Foundation. SVG acknowledges funding support from NSERC.

## References

1. Dian C, Perez-Dorado I, Riviere F, Asensio T, Legrand P, Ritzefeld M, Shen M, Cota E, Meinnel T, Tate EW et al: High-resolution snapshots of human N-myristoyltransferase in action illuminate a mechanism promoting N-terminal Lys and Gly myristoylation. Nat Commun 2020, 11(1): 1132.

2. Chauhan R, Datzkiw D, Varma Shrivastav S, Shrivastav A: In silico identification of microRNAs predicted to regulate N-myristoyltransferase and Methionine Aminopeptidase 2 functions in cancer and infectious diseases. PLoS One 2018, 13(3):e0194612.

3. McIlhinney RA, McGlone K, Willis AC: Purification and partial sequencing of myristoyl-CoA:protein N-myristoyltransferase from bovine brain. Biochem J 1993, 290 (Pt 2):405–410.

4. Podell S, Gribskov M: Predicting N-terminal myristoylation sites in plant proteins. BMC Genomics 2004, 5(1):37.

5. Galvin BD, Li Z, Villemaine E, Poole CB, Chapman MS, Pollastri MP, Wyatt PG, Carlow CK: A target repurposing approach identifies N-myristoyltransferase as a new candidate drug target in filarial nematodes. PLoS Negl Trop Dis 2014, 8(9):e3145.

6. Fang W, Robinson DA, Raimi OG, Blair DE, Harrison JR, Lockhart DE, Torrie LS, Ruda GF, Wyatt PG, Gilbert IH et al: N-myristoyltransferase is a cell wall target in Aspergillus fumigatus. ACS Chem Biol 2015, 10(6):1425–1434.

7. Corpas-Lopez V, Moniz S, Thomas M, Wall RJ, Torrie LS, Zander-Dinse D, Tinti M, Brand S, Stojanovski L, Manthri S et al: Pharmacological Validation of N-Myristoyltransferase as a Drug Target in Leishmania donovani. ACS Infect Dis 2019, 5(1):111–122.

8. Towler DA, Adams SP, Eubanks SR, Towery DS, Jackson-Machelski E, Glaser L, Gordon JI: Purification and characterization of yeast myristoyl CoA:protein N-myristoyltransferase. Proc Natl Acad Sci U S A 1987, 84(9):2708–2712.

9. Shrivastav A, Suri SS, Mohr R, Janardhan KS, Sharma RK, Singh B: Expression and activity of N-myristoyltransferase in lung inflammation of cattle and its role in neutrophil apoptosis. Vet Res 2010, 41(1):9.

10. Duronio RJ, Knoll LJ, Gordon JI: Isolation of a Saccharomyces cerevisiae long chain fatty acyl:CoA synthetase gene (FAA1) and assessment of its role in protein N-myristoylation. J Cell Biol 1992, 117(3):515–529.

11. Udenwobele DI, Su RC, Good SV, Ball TB, Varma Shrivastav S, Shrivastav A: Myristoylation: An Important Protein Modification in the Immune Response. Front Immunol 2017, 8:751.

12. Utsumi T, Matsuzaki K, Kiwado A, Tanikawa A, Kikkawa Y, Hosokawa T, Otsuka A, Iuchi Y, Kobuchi H, Moriya K: Identification and characterization of protein N-myristoylation occurring on four human mitochondrial proteins, SAMM50, TOMM40, MIC19, and MIC25. PLoS One 2018, 13(11):e0206355.

13. Giang DK, Cravatt BF: A second mammalian N-myristoyltransferase. J Biol Chem 1998, 273(12):6595–6598.

14. Kumar S, Sharma RK: N-terminal region of the catalytic domain of human N-myristoyltransferase 1 acts as an inhibitory module. PLoS One 2015, 10(5):e0127661.

15. Yang SH, Shrivastav A, Kosinski C, Sharma RK, Chen MH, Berthiaume LG, Peters LL, Chuang PT, Young SG, Bergo MO: N-myristoyltransferase 1 is essential in early mouse development. J Biol Chem 2005, 280(19):18990–18995.

16. Ducker CE, Upson JJ, French KJ, Smith CD: Two N-myristoyltransferase isozymes play unique roles in protein myristoylation, proliferation, and apoptosis. Mol Cancer Res 2005, 3(8):463–476.

17. Thinon E, Morales-Sanfrutos J, Mann DJ, Tate EW: N-Myristoyltransferase Inhibition Induces ER-Stress, Cell Cycle Arrest, and Apoptosis in Cancer Cells. ACS Chem Biol 2016, 11(8):2165–2176.

18. Shrivastav A, Varma S, Saxena A, DeCoteau J, Sharma RK: N-myristoyltransferase: a potential novel diagnostic marker for colon cancer. J Transl Med 2007, 5:58.

19. Selvakumar P, Lakshmikuttyamma A, Shrivastav A, Das SB, Dimmock JR, Sharma RK: Potential role of N-myristoyltransferase in cancer. Prog Lipid Res 2007, 46(1):1–36.

20. Jacquier M, Kuriakose S, Bhardwaj A, Zhang Y, Shrivastav A, Portet S, Varma Shrivastav S: Investigation of Novel Regulation of N-myristoyltransferase by Mammalian Target of Rapamycin in Breast Cancer Cells. Sci Rep 2018, 8(1):12969.

21. Clegg RA, Gordge PC, Miller WR: Expression of enzymes of covalent protein modification during regulated and dysregulated proliferation of mammary epithelial cells: PKA, PKC and NMT. Adv Enzyme Regul 1999, 39:175–203.

22. Mousnier A, Bell AS, Swieboda DP, Morales-Sanfrutos J, Perez-Dorado I, Brannigan JA, Newman J, Ritzefeld M, Hutton JA, Guedan A et al: Fragment-derived inhibitors of human N-myristoyltransferase block capsid assembly and replication of the common cold virus. Nat Chem 2018, 10(6):599–606.

23. Seaton KE, Smith CD: N-Myristoyltransferase isozymes exhibit differential specificity for human immunodeficiency virus type 1 Gag and Nef. J Gen Virol 2008, 89(Pt 1):288–296.

24. Rozewicki J, Li S, Amada KM, Standley DM, Katoh K: MAFFT-DASH: integrated protein sequence and structural alignment. Nucleic Acids Res 2019, 47(W1):W5–W10.

25. Kumar S, Stecher G, Li M, Knyaz C, Tamura K: Mega X: Molecular Evolutionary Genetics Analysis across Computing Platforms. Mol Biol Evol 2018, 35(6):1547–1549.

26. Yu G, Lam TT, Zhu H, Guan Y: Two Methods for Mapping and Visualizing Associated Data on Phylogeny Using Ggtree. Mol Biol Evol 2018, 35(12):3041–3043.

27. Yegorov S, Good S: Using paleogenomics to study the evolution of gene families: origin and duplication history of the relaxin family hormones and their receptors. PLoS One 2012, 7(3):e32923.

28. Sacerdot C, Louis A, Bon C, Berthelot C, Roest Crollius H: Chromosome evolution at the origin of the ancestral vertebrate genome. Genome Biol 2018, 19(1):166.

29. Nguyen Ba AN, Pogoutse A, Provart N, Moses AM: NLStradamus: a simple Hidden Markov Model for nuclear localization signal prediction. BMC Bioinformatics 2009, 10:202.

30. http://nls-mapper.iab.keio.ac.jp/cgi-bin/NLS_Mapper_help.cgi

31. Brameier M, Krings A, MacCallum RM: NucPred--predicting nuclear localization of proteins. Bioinformatics 2007, 23(9): 1159–1160.

32. Opanasopit P, Rojanarata T, Apirakaramwong A, Ngawhirunpat T, Ruktanonchai U: Nuclear localization signal peptides enhance transfection efficiency of chitosan/DNA complexes. Int J Pharm 2009, 382(1-2):291–295.

33. Fang Y, Jang HS, Watson GW, Wellappili DP, Tyler BM: Distinctive Nuclear Localization Signals in the Oomycete Phytophthora sojae. Front Microbiol 2017, 8:10.

34. Strom AC, Weis K: Importin-beta-like nuclear transport receptors. Genome Biol 2001, 2(6):REVIEWS3008.

35. Sidorova JM, Mikesell GE, Breeden LL: Cell cycle-regulated phosphorylation of Swi6 controls its nuclear localization. Mol Biol Cell 1995, 6(12):1641–1658.

36. Johnson LN, Barford D: The effects of phosphorylation on the structure and function of proteins. Annu Rev Biophys Biomol Struct 1993, 22:199–232.

37. Schnell JD, Hicke L: Non-traditional functions of ubiquitin and ubiquitin-binding proteins. J Biol Chem 2003, 278(38):35857–35860.

38. Lin YY, Lu JY, Zhang J, Walter W, Dang W, Wan J, Tao SC, Qian J, Zhao Y, Boeke JD et al: Protein acetylation microarray reveals that NuA4 controls key metabolic target regulating gluconeogenesis. Cell 2009, 136(6):1073–1084.

39. Vlastaridis P, Kyriakidou P, Chaliotis A, Van de Peer Y, Oliver SG, Amoutzias GD: Estimating the total number of phosphoproteins and phosphorylation sites in eukaryotic proteomes. Gigascience 2017, 6(2):1–11.

40. Zhang F, White RL, Neufeld KL: Phosphorylation near nuclear localization signal regulates nuclear import of adenomatous polyposis coli protein. Proc Natl Acad Sci U S A 2000, 97(23):12577–12582.

41. Smith KM, Di Antonio V, Bellucci L, Thomas DR, Caporuscio F, Ciccarese F, Ghassabian H, Wagstaff KM, Forwood JK, Jans DA et al: Contribution of the residue at position 4 within classical nuclear localization signals to modulating interaction with importins and nuclear targeting. Biochim Biophys Acta Mol Cell Res 2018, 1865(8):1114–1129.

42. Lange A, McLane LM, Mills RE, Devine SE, Corbett AH: Expanding the definition of the classical bipartite nuclear localization signal. Traffic 2010, 11(3):311–323.

43. Lange A, Mills RE, Devine SE, Corbett AH: A PY-NLS nuclear targeting signal is required for nuclear localization and function of the Saccharomyces cerevisiae mRNA-binding protein Hrp1. J Biol Chem 2008, 283(19):12926–12934.

44. Kosugi S, Hasebe M, Matsumura N, Takashima H, Miyamoto-Sato E, Tomita M, Yanagawa H: Six classes of nuclear localization signals specific to different binding grooves of importin alpha. J Biol Chem 2009, 284(1):478–485.

45. Nakada R, Hirano H, Matsuura Y: Structure of importin-alpha bound to a non-classical nuclear localization signal of the influenza A virus nucleoprotein. Sci Rep 2015, 5:15055.

