## Supplementary figures and images for "N-myristoyltransferase: Tracing Steps Backwards to Find a Way Forward"

### Vertebrate Tree

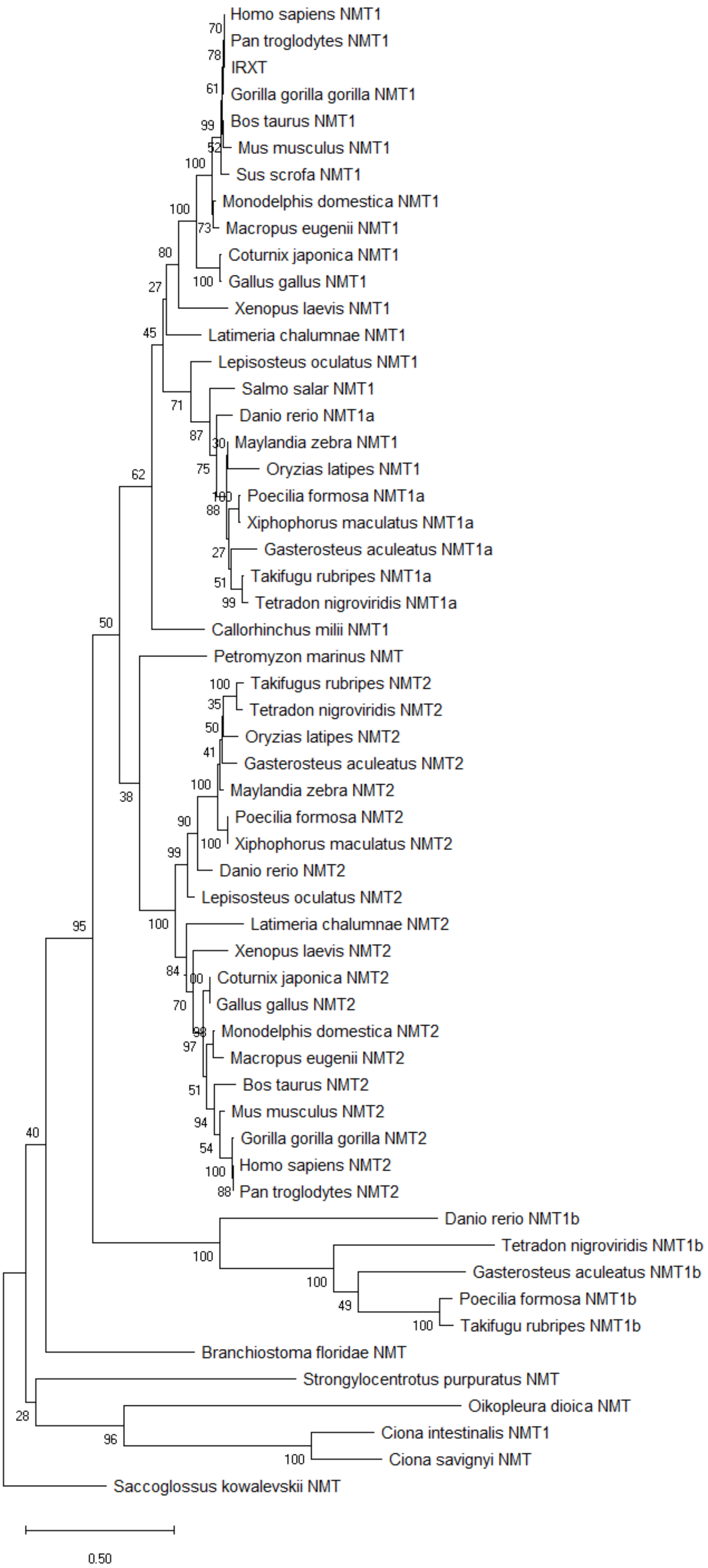
